# International spread of emerging multidrug-resistant *Rhodococcus equi*

**DOI:** 10.1101/2022.02.15.480578

**Authors:** Jorge Val-Calvo, Jane Darcy, James Gibbons, Alan Creighton, Claire Egan, Thomas Buckley, Achim Schmalenberger, Ursula Fogarty, Mariela Scortti, José A. Vázquez-Boland

## Abstract

The recently characterized multidrug-resistant clone of the animal and human zoonotic pathogen *Rhodococcus equi*, MDR-RE 2287, has been circulating among horse farms in the United States (US) since the early 2000’s. Here, we report the first documented detection of the MDR-RE 2287 clone outside the US. The finding highlights the risk of MDR-RE spreading internationally with the movement of horses.

## Introduction

*Rhodococcus equi* is a soil-borne aerobic actinomycete that causes pyogranulomatous infections in animals and people. Human infections are opportunistic, can be linked to exposure to farm environments, and are zoonotic in origin (1-3). Although clinical *R. equi* infections are relatively rare in most animal species, foals are commonly affected and develop a potentially life-threatening disease characterized by purulent pneumonia, with a high incidence in equine farms worldwide (4). The mainstay treatment of foal rhodococcosis consists in long courses of a macrolide and rifampin. Systematically applied since the 1980’s, no significant resistance was detected until the early 2000’s following mass prophylactic application of the combination therapy at endemic farms in the United States (US) (5, 6). The emerging dual macrolide-rifampin resistance was attributable to a multidrug-resistant *R. equi* (MDR-RE) clone, named *“*2287”, which has been spreading among horse farms across the US. MDR-RE 2287 arose by co-acquisition of the conjugative plasmid pRErm46 and a specific *rpoB*^S531F^ (TCG→TTC) mutation conferring high-level rifampin resistance (7, 8). pRErm46 specifies resistance to macrolides, lincosamides and streptogramins via the *erm*(46) gene carried on Tn*RErm46*, a highly mobile transposon, and to sulfonamides, streptomycin, spectinomycin, tetracycline and doxycycline via a class 1 integron (C1I) and associated *tetRA* determinant (9). Only detected so far in the US, MDR-RE was predicted to disseminate to other countries with the movement of equines (10).

## The study

Following the characterization of MDR-RE in 2019 (8), we established an informal surveillance network with colleagues in North and South America, Europe, UK, Africa, Asia and Australia. Collaborating laboratories were asked to review their retrospective *R. equi* collections and prospectively identify any isolate with a minimum inhibitory concentration (MIC) for erythromycin ≥4 µg/ml potentially denoting *erm*(46)-mediated macrolide resistance. Two equine clinical strains from necropsied foals in Ireland met the criterion: PAM 2528 recovered in 2016 and PAM 2578 in 2021. Both originated from the same farm and had MICs of ≥32 µg/ml for erythromycin and >256 µg/ml for rifampin, consistent with MDR-RE’s resistance phenotype (7-10). No other macrolide-resistant *R. equi* strains were notified by our collaborators to date.

Both isolates were confirmed as *erm*(46)-positive by PCR and to carry the *rpoB*^S531F^ mutation unique to the MDR-RE 2287 clone using previously described methods (9). A PCR designed to detect C1I-*tetRA* deletions in pRErm46 (9) (oligonucleotides C1I-check-F 5’-ccgagatgtgtcggacttc and C1I-check-R 5’-cgccgaagaacaacccgaggatg), observed in a proportion of recent MDR-RE isolates (9,10), showed the resistance plasmid was of the ΔC1I-*tetRA* type. Accordingly, PAM 2528 and PAM 2578 were susceptible to trimethoprim-sulfamethoxazole, streptomycin, spectinomycin, and tetracycline, which the pRErm46 C1I-*tetRA* determinant confers resistance to (9).

Genomic DNA was extracted and paired-end Illumina sequenced, and reads were quality-checked using FastQC v0.11.9 (https://www.bioinformatics.babraham.ac.uk/projects/fastqc/), trimmed using TrimmomaticPE v0.39 (11), and assembled using SPAdes v3.15.2 (12). Presence of pRErm46 sequences in the draft genomes was confirmed using BlastN. ParSNP v1.5.6 and FastTree were used to build approximate-maximum likelihood (ML) trees based on core single nucleotide polymorphisms (SNPs) (13) to determine the position of the isolates in the *R. equi* population structure. The output *R. equi* tree showed that the two macrolide-and rifampin-resistant isolates from Ireland belonged to the MDR-RE 2287 clone (Figure 1).

**Figure 1.**
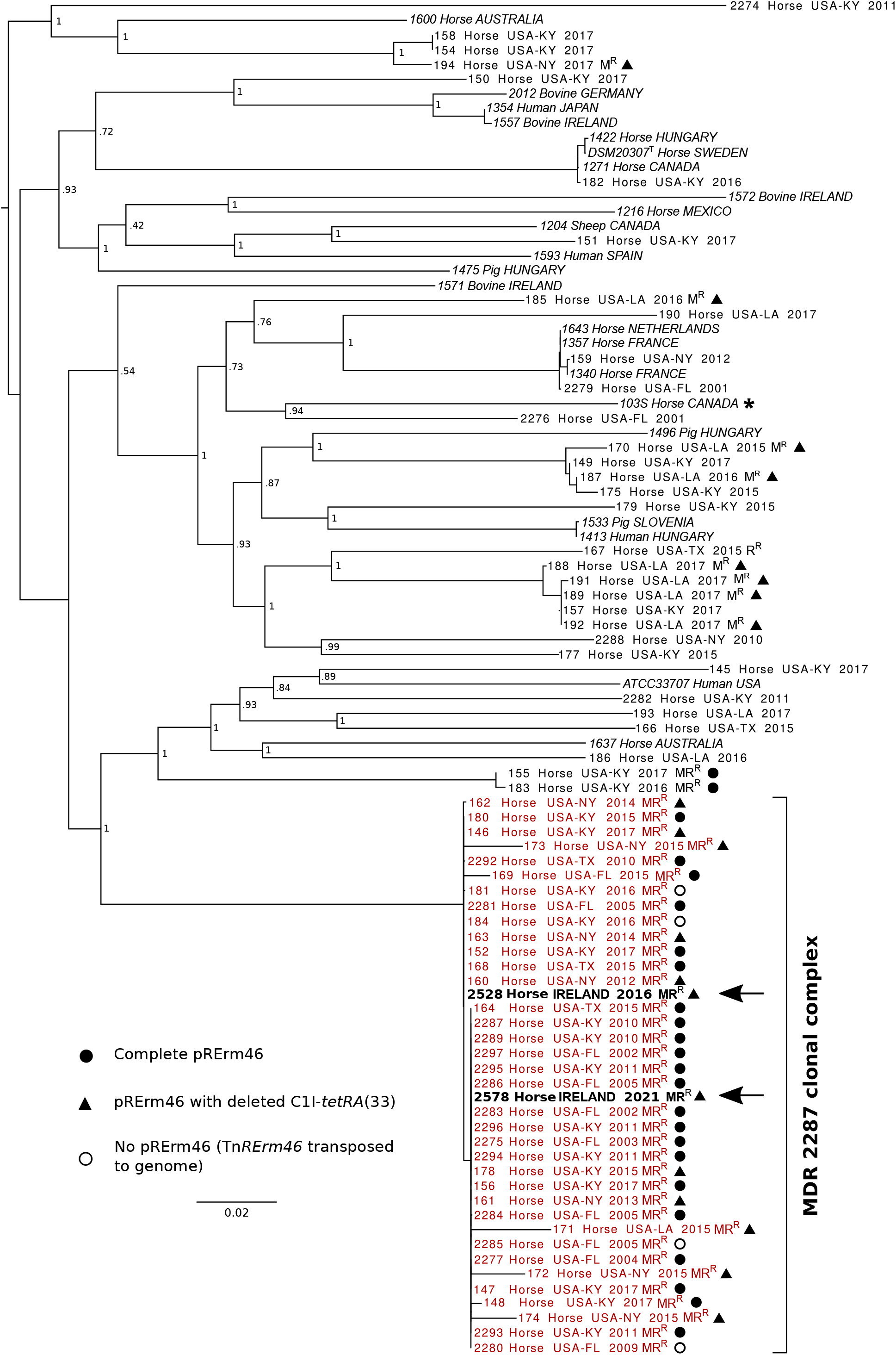
*R. equi* whole-genome phylogenetic analysis identifies two equine isolates from Ireland (arrows) as members of the MDR-RE 2287 clonal complex. Tree constructed with ParSNP in the Harvest suite using the complete genome of *R. equi* 103S as a reference (indicated with an asterisk; GenBank accession no. FN563149). Analysis performed using 92 *R. equi* genome sequences including 22 from a previously reported *R. equi* diversity set (15) (labeled in italics) and a collection of 68 macrolide-resistant and -susceptible equine isolates from the US where MDR-RE is currently circulating (8, 10) (in regular font). The latter include 36 MDR-RE 2287 isolates (highlighted in red, the Irish strains in black and indicated by arrows), 10 isolates representing spillages of the pRErm46 plasmid to other *R. equi* genotypes, and 22 control susceptible isolates (8, 10). Labels indicate geographical origin, year of isolation and resistance phenotype when applicable (MR^R^, macrolide and rifampin; M^R^, only macrolides; R^R^, only rifampin). pRErm46 carriage in macrolide-resistant isolates is indicated by symbols (see inset legend). The empty circles indicate MDR-RE isolates where pRErm46 has been lost after transposition of the Tn*RErm46* element to the host genome (8). Numbers in the nodes indicate bootstrap values for 1,000 replicates. Tree graphed with FigTree (http://tree.bio.ed.ac.uk/software/figtree).

Since the short genetic distances compressed the branching of the MDR-RE 2287 isolates, to explore in more detail their relationships, we repeated the phylogenetic analysis with only the clonal genomes (Figure 2). For this analysis, core SNPs were detected using SNIPPY (v4.6.0, https://github.com/tseemann/snippy), which we found avoids genome alignment errors observed with ParSNP that significantly distort the phylogenetic reconstruction of virtually identical isolates (average of 58 SNPs between MDR-RE 2287 isolates vs 29,743±3,798 for random *R. equi* strains).

**Figure 2.**
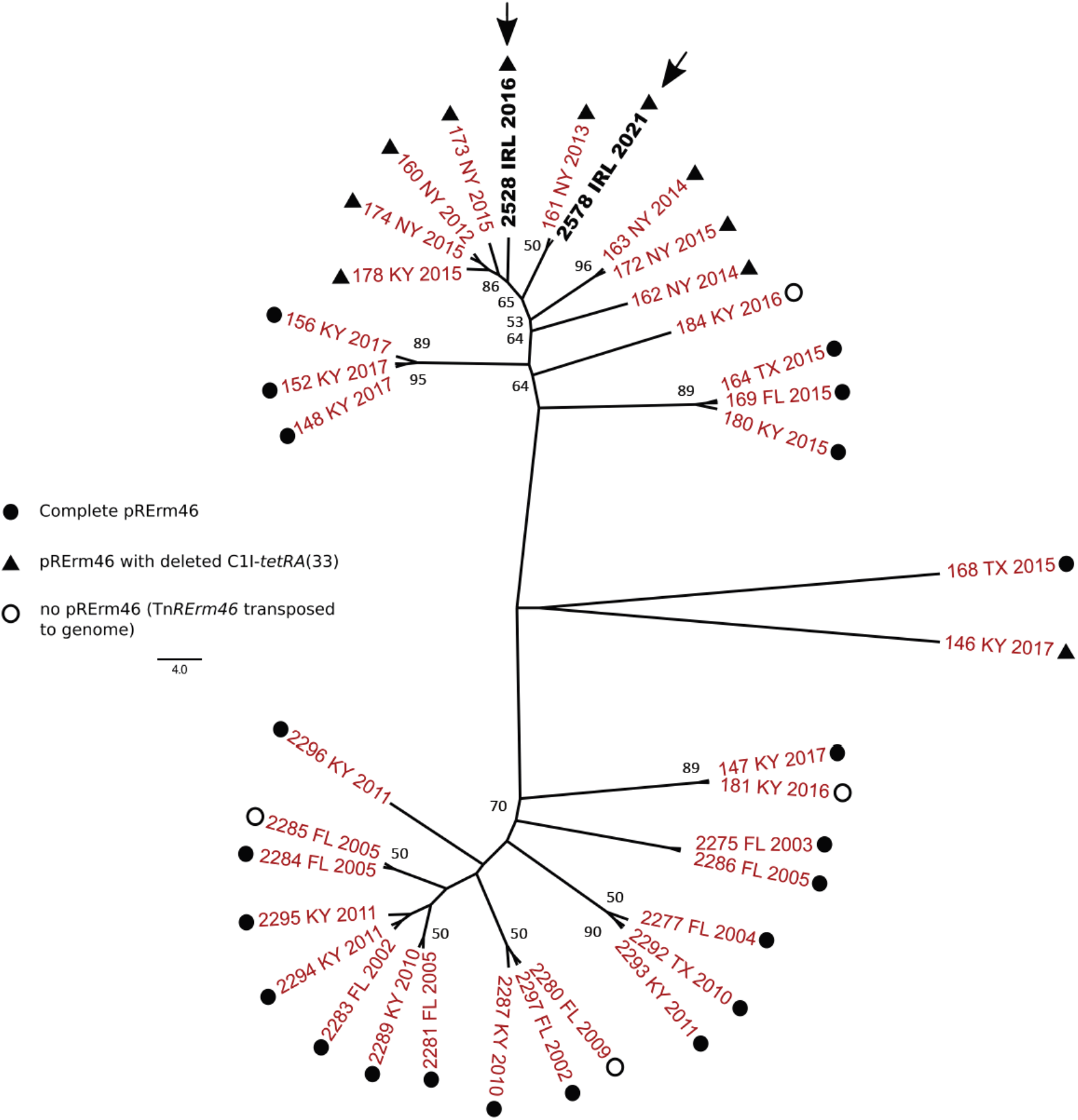
Unrooted ML tree of MDR-RE 2287 clonal complex showing the relationships of the isolates from Ireland (in black and indicated by arrows). Whole-genome phylogeny inferred from 45 parsimony informative sites using SNIPPY and IQtree for tree recontruction. The genome of the prototype MDR-RE 2287 isolate PAM 2287 (NCBI assembly accession no. GCA_002094405.1) was used as a reference for SNP calling. Best-Fit model selected by ModelFinder module was K3Pu+F+ASC. Bootstrap values ≥50 are shown. Geographical source: FL, Florida; IRL, Ireland, KY, Kentucky; NY, New York; TX, Texas. pRErm46 type is indicated by symbols (see inset legend). Tree drawn with FigTree (http://tree.bio.ed.ac.uk/software/figtree).

The consensus ML tree subdivided the MDR-RE 2287 clonal complex in two main sublineages comprising respectively older (2002 to 2011) and younger (2015 onwards) isolates. PAM 2528 and PAM 2578 clustered close to each other at the top branches of the younger sublineage together with all the isolates from New York in the analyzed MDR-RE 2287 collection, suggesting a common source origin. This was further supported by finding that both PAM 2528 and PAM 2578 carried ΔC1I-*tetRA* pRErm46 variants, like all New York isolates (Figure 2).

## Conclusions

We report the first documented international spread of the MDR-RE 2287 clone that has been circulating in the US since the 2000’s (8, 10). *R. equi* 2287 appears to be following the same pattern of the pandemic MDR clones of human bacterial pathogens, which within a few years after emergence and initial local expansion become globally disseminated (14). This is taking place at a much slower pace with MDR-RE, likely because of the lesser opportunities for transmission afforded by horse trade and inter-horse contacts compared to human interactions and travel.

While the analyzed MDR-RE 2287 isolates were from various US origins, the two Irish strains clustered together in the tree with (all seven) isolates from New York. The latter were recovered over a period of several years since 2012, pointing to an MDR 2287 subpopulation established in a farm(s) in New York State as the likely source of the isolates recovered in Ireland. The “New York” MDR-RE 2287 subpopulation is characterized by possessing the ΔC1I-*tetRA* pRErm46 plasmid variant, also carried by the Irish isolates but only exceptionally by other members of the MDR-RE 2287 complex (Figures 1 and 2).

The positioning of the Irish isolates in two separate sub-branches of the “New York” radiation (Figure 2) may indicate they represent independent, temporally distinct import events that took place around 2016 and 2021 involving different subclones of that particular subpopulation. That MDR-RE 2287 was not detected again in Ireland until five years later, either from repeated environmental samples collected on the affected farm or routine screening of equine *R. equi* clinical isolates, is a possible indication that it might not have persisted after its first appearance in 2016. This scenario may be explained by the different *R. equi*-targeted equine farm management practices in Ireland compared to the US, where application of mass antibioprophylaxis likely favored the emergence, maintenance and spread of MDR-RE (6, 8). However, the genetic distance between the two Irish isolates is comparable to that between the seven New York isolates, recovered from 2012 to 2015. It cannot therefore be excluded that PAM 2528 and PAM 2578 represent successive isolations of a locally evolving single imported subclone (21 SNPs difference over 5 M bp ≈ 1×10^−6^ substitutions per site per year, consistent with normal genetic drift values).

A Kentucky isolate of the ΔC1I-*tetRA* type was also located in a terminal branch of the “New York” cluster, whereas Kentucky isolates with complete pRErm46 plasmids were positioned at basal bifurcations of the radiation (e.g. the 148, 152 and 153 cluster) (Figure 2). This suggests a transmission history in which a relatively recent MDR-RE 2287 subclone which acquired at some point a ΔC1I-*tetRA* deletion, possibly originating from Kentucky (where MDR-RE emerged and is prevalent) (8, 9), became endemic in a New York farm(s) and was subsequently transferred, either directly or indirectly, to Ireland. International trade in thoroughbred horses is frequent and the affected farm in Ireland received horses from America, Europe, UK and other Irish farms on a regular basis. Previous phylogenomic studies provided evidence of global circulation of *R. equi* genomotypes, probably linked to livestock trade (15). Our findings here consolidate this notion and warn about the risk for MDR-RE becoming internationally disseminated over time with horse movements.

It is worth noting that our study is not comprehensive but based on the voluntary collaboration of a restricted number of international colleagues, and thus MDR-RE may have also spread to other countries. It would be important to actively monitor the occurrence of the emerging MDR-RE 2287 clone for which, as our data highlight, *erm*(46) and the *rpoB*^S531F^ (TCG→TTC) mutation can be used as molecular markers, eventually complemented with pRErm46/ΔC1I-*tetRA* variant detection.

## Acknowledgements

We are grateful to the network of international collaborators for their participation in macrolide-and rifampin-resistant *R. equi* monitoring.

Work on MDR *R. equi* at JV-B’s laboratory is supported by the Horserace Betting Levy Board (HBLB grant prj-796).

## Notes

### Competing Interest Statement

The authors have declared no competing interest.

